# DEGAP: Dynamic Elongation of a Genome Assembly Path

**DOI:** 10.1101/2023.04.25.538224

**Authors:** Yicheng Huang, Ziyuan Wang, Monica A. Schmidt, Jianwei Zhang

## Abstract

Genome assembly remains to be a major task in genomic research. Despite the development over the past decades of different assembler software programs and algorithms, it is still a great challenge to assemble a complete genome with no gaps. With the latest DNA Circular Consensus Sequencing (CCS) technology, several assembler programs now can build a genome from raw sequencing data to contigs, however, some complex sequence regions remain as unresolved gaps. Here, we present a novel gap-filling software DEGAP that can resolve gap regions in genomes by utilizing the dual advantages of accuracy and length of high-fidelity (HiFi) reads. DEGAP optimizes HiFi reads by identifying the differences between reads and provides ‘GapFiller’ or ‘CtgLinker’ modes to eliminate or shorten gaps in genomes. DEGAP adopts a cyclic elongation strategy that automatically and dynamically adjusts parameters according to the complexity of the sequences and selects the optimal extension path. DEGAP has already been successfully applied to decipher complex genomic regions in several projects and may be widely employed to generate more gap-free genomes.

## Introduction

DNA sequencing had a long and inspiring history of development. Each generation of sequencing technologies has propelled the decrypting of the genetic code and understanding of the biology and evolution of species. Complete and accurate genome assembly is an essential component for genomic analyses and historically has relied on long sequencing reads that are often inaccurate, and on accurate short reads. Pacific Biosciences (PacBio) currently provides high-fidelity (HiFi) sequencing mode to increase both the accuracy and the length of sequence reads, routinely producing over 10 kbp long reads with 99.9% accuracy^(1)^.

With the development of sequencing technologies, several software tools (e.g., Canu^(2)^, MECAT^(3)^, FALCON^(4)^, Flye^(5)^, hifiasm^(6)^, Wtdbg^(7)^) have become available for *de novo* assembling genomes by using various raw sequencing data, which in many ways greatly accelerates the genome assembly process. However, gaps still remain in an assembly due to either low sequencing depth or high sequence complexity. Hence, a high-resolution tool capable of easily and accurately closing gaps is a necessary requirement to assemble complete genome sequences.

In this study, we developed an automatic tool, named DEGAP (Dynamic Elongation of a Genome Assembly Path), to elongate the edge sequences between gap areas in the genome and find a continuous path to extend edge sequences with HiFi data. DEGAP can efficiently resolve gaps in non-chromosome-level assemblies without generating more sequencing data and cost-effectively reduces the overall gap number in an entire genome. DEGAP’s ability to resolve gap regions will undoubtedly make it widely useful in the generation of more gap-free genomes.

## Results

### Two gap-closure modes: GapFiller and CtgLinker

DEGAP provides GapFiller and CtgLinker modes for users to close gaps in *de novo* assembled genomes by elongating the edge sequences of gap regions, which may fit two scenarios for different genome assemblies. The GapFiller mode focuses on filling known gaps specifically in chromosome-level assemblies: DEGAP takes both left and right sequences of gap regions as seed sequences and uses raw HiFi reads to incrementally elongate the seed sequences to eventually reach the sequences on the other side of each gap. The CtgLinker mode is designed for handling non-chromosome-level assemblies of unknown gaps: In this mode, DEGAP first filters out the overly short contigs, then cut off the edges of the contigs to reduce the mis-assembly caused by sequencing errors and pursue accurate elongation results. DEGAP elongates all contigs with supplied HiFi data, assesses the potentially neighbored contigs one-by-one in the dataset by their overlaps, then connects them if the elongated sequence finds a well-matched contig. The automatic process continually performs elongation tasks until all gaps are filled or no extension sequences (or reads) found.

### Dynamic process of assembly path elongation

DEGAP adopts a cyclic elongation strategy that automatically uses dynamic parameter thresholds to select sequences for extension according to the complexity of the actual situation, which can be divided into 4 steps: (i) extracting reads which are located at one side of the gap, (ii) processing the candidate extension reads or sequences, (iii) elongating the edge sequence using the optimal candidate in the previous step, (iv) determining if the elongated sequence reaches the other side of the gap (Fig. 1a). In the fourth step, DEGAP checks if the elongated sequence has reached the other side of the gap to determine to stop or continue the elongation process using the elongated sequences as the input sequence for the next elongation round (Fig. 1b).

**Fig. 1:**
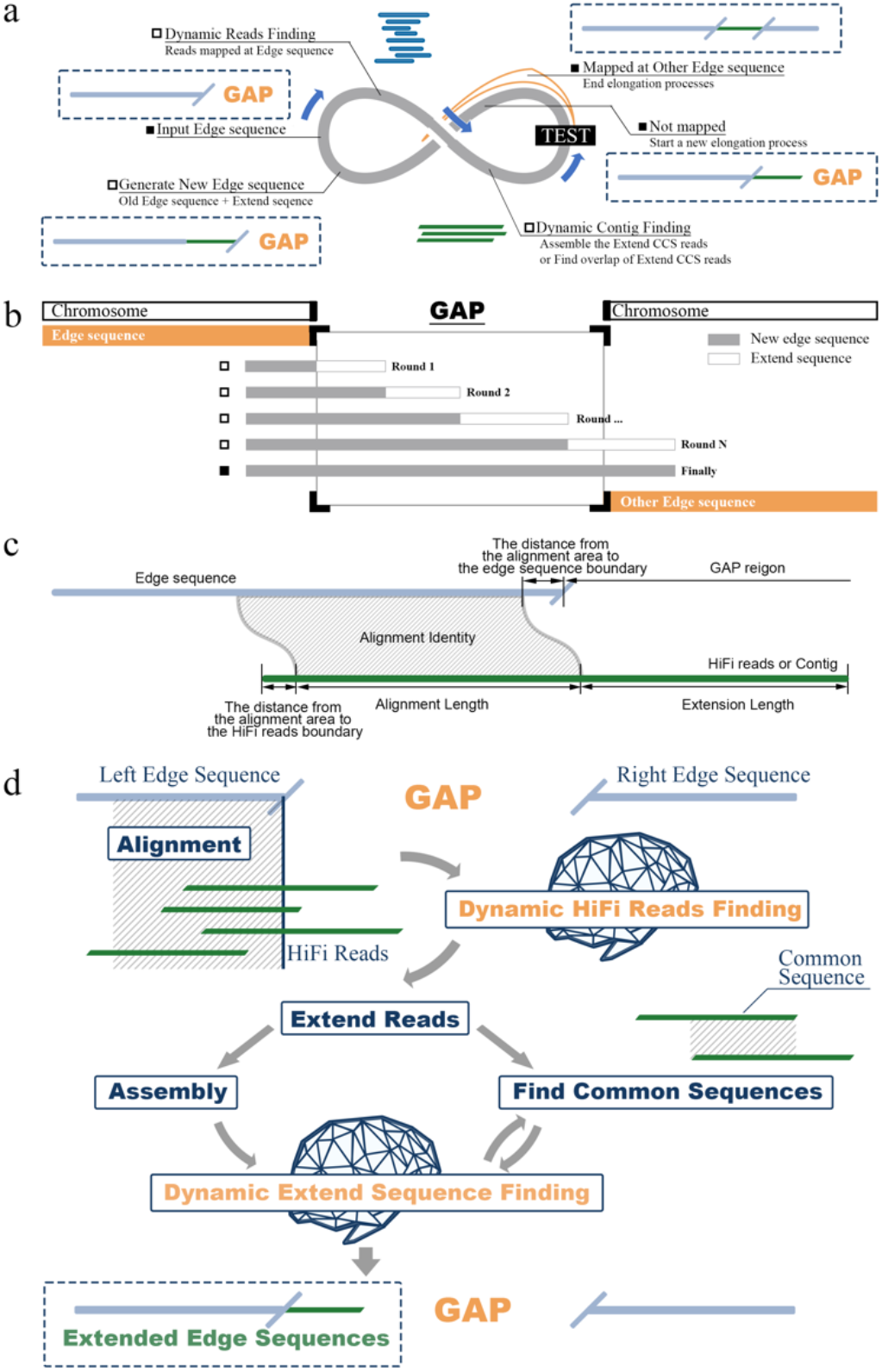
DEGAP pipeline. **a**, Cyclic elongation strategy of DEGAP. **b**, Complete elongation process when DEGAP filled the gaps. **c**, Parameters to evaluate whether HiFi reads / sequences are eligible to extend the edge sequence. **d**, Two core dynamic process of DEGAP.

In each round of elongation, the basic principle for DEGAP is to select the correct HiFi reads or extension sequence to elongate the edge sequences between gaps. The accuracy of assessing whether extension HiFi reads/sequence are eligible for elongation depends on the alignments to edge sequence. Considering the differences in the input edge sequence for each round, DEGAP will dynamically change threshold values of four different parameters to select the alignment blocks: (i) the distance from the alignment area to the sequence boundary, (ii) alignment identity, (iii) alignment length and, (iv) extension length of the reads (Fig. 1c, Supplement Fig. 1, 2).

During the elongation process for each round, DEGAP applies a dynamic decision-making rule to find and select extension reads or contigs in consideration of both assembly error and qualified alignment results (Fig. 1d). Using minimap2^(8)^ to map all HiFi reads back to one edge of a gap, the dynamic reads finding process in DEGAP finds a set of candidate extension reads that have a high identity and high coverage alignment with the edge for potentially extending the edge sequence (Fig. 1d, Supplementary Fig. 1). Subsequently, the dynamic extension sequence finding process may *de novo* assemble the extension reads obtained in the previous step by using hifiasm^(6)^ or find the common sequences between extension HiFi reads, and then pick up the best extension sequence to precisely elongate the edge sequence (Fig. 1d, Supplementary Fig. 2).

DEGAP determines qualified alignment results by considering both maximum alignment identity and maximum alignment length against edge sequence (Supplementary Fig. 3). Meanwhile, DEGAP also produces a fuzzy alignment result in second dynamic process when no alignment results (i.e., repeat-caused multiple alignment results) are qualified (Supplementary Fig. 3d, e).

By taking the edge sequence at one end of the gap as a seed, DEGAP generates a continuous sequence path through the cyclic extension process. Therefore, there are three possible outcomes of the DEGAP process: (i) the gap is filled; (ii) no extension reads/sequence found; (iii) no extension reads found and the process stuck in an infinite loop.

### Two study cases of DEGAP

#### 1) MH63 centromere sequences resolved in the GapFiller mode

Centromeric DNA usually contains highly repetitive DNAs, such as satellite DNAs, which become one of the difficulties in genome assembly. In this study, we used the *Oryza sativa* gap-free genome assembled with multiple sequencing data as a reference, which contains complete centromere sequences including *CentO* satellite repeats^(9, 10)^. By using DEGAP, we bidirectionally reproduced all centromere sequences in MH63RS3 and compared them with the corresponding sequences in MH63RS3 to validate the DEGAP results (Fig. 2). As the result, DEGAP could fill the complete centromere sequences in Chr03, Chr06, Chr10 from both directions, and Chr07, Chr09 from the left side, and Chr12 from the right side of centromere (Fig. 2a), including those known gaps on Chr01, Chr02, Chr07, Chr09, Chr11 in MH63RS2.

**Fig. 2:**
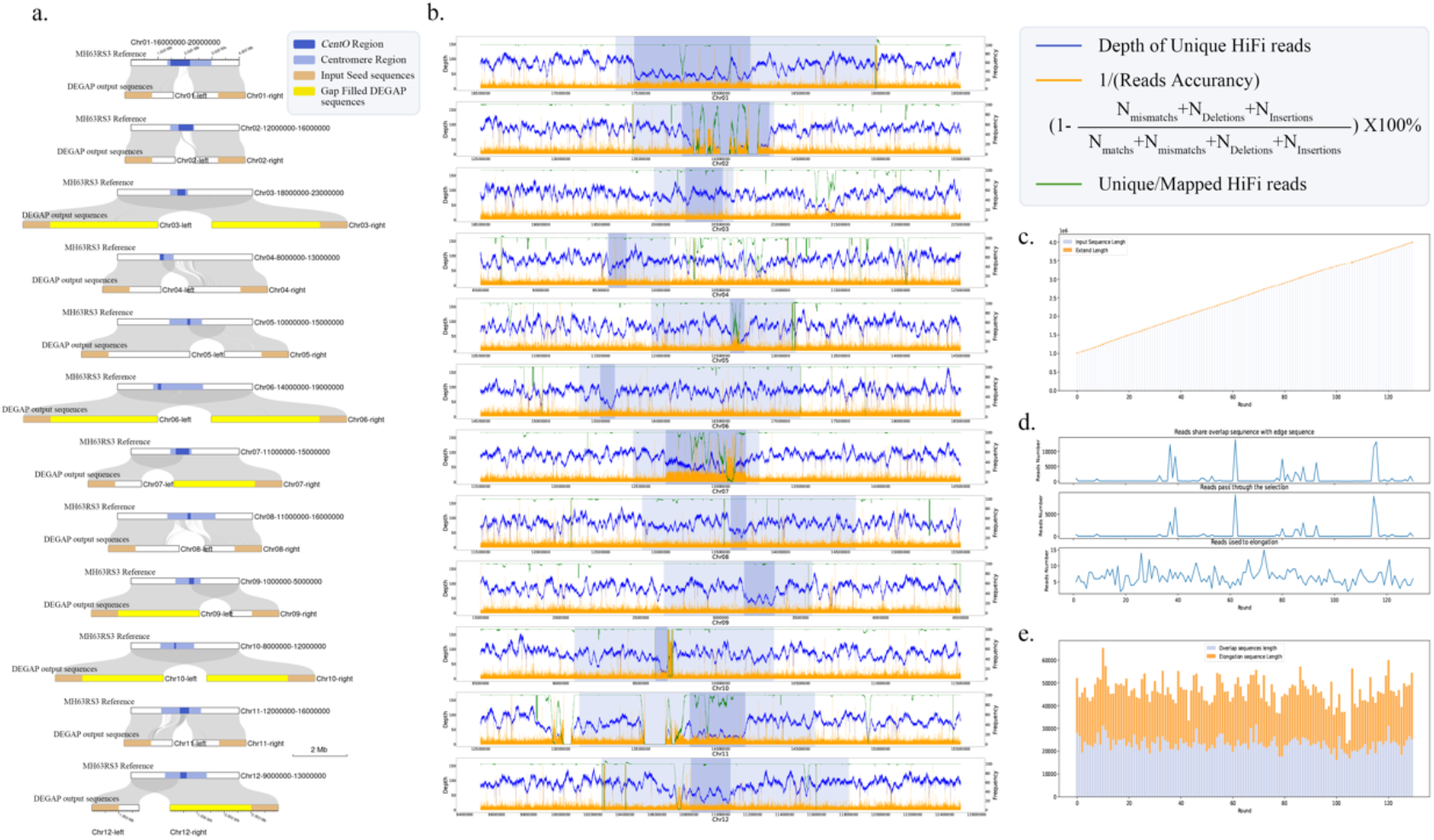
DEGAP-resolved Results of MH63 centromere sequences in GapFiller mode. **a**, Alignment of DEGAP results with the MH63 reference genome. Chr03, Chr06, Chr07, Chr09, Chr10 and Chr12 of MH63 have complete centromeric sequences elongated by DEGAP in GapFiller mode. Dark blue represents *CentO* satellite repeats. Light Blue represents centromere sequences. Brown represents DEGAP input sequences. Yellow represents DEGAP-elongated entire centromere regions. **b**, Sequences complexity in the MH63RS3 centromere sequences. The closer to the centromere region, the lower the depth of unique HiFi reads, the lower the accuracy of reads, the more difficult it is for DEGAP to obtain correct reads. Blue line represents the unique HiFi read depth value. Yellow line represents the reads inaccuracy. The definition that we choose is the proportion of base mismatch, deletion and insertion in the matching columns in the alignment of the reads to the reference sequence. Green line indicates the ratio of unique HiFi reads among all mapped HiFi reads in the same area. We use this to assess the difficulty for DEGAP to obtain correct reads. **c**, The input sequences length and extension length in each round of Chr03 left side DEGAP elongation. **d**, The number of reads matched edge sequences, reads passed through selection and reads used to elongation in each round of Chr03 left side DEGAP elongation. **e**, The length of overlap and elongated sequences in elongation sequences in each round of Chr03 left side of DEGAP elongation.

#### 2) Human genome sequences improved with the CtgLinker mode

The first near-complete human reference genome (GRCh38.p13) was released in 2013^(11)^ by the Genome Reference Consortium, which leaves 349 unfinished repetitive and polymorphic regions as gaps. A Telomere-to-Telomere (T2T) version (CHM13) of the reference human genome was assembled in 2022 by using PacBio HiFi and Oxford Nanopore ultralong-read sequencing technologies^(12)^. The T2T-CHM13 reference genome (CHM13v1.1) was used to test the DEGAP performance and the former non-T2T assembly (CHM13v0.7) can be used as input dataset to test if DEGAP can solve non-chromosome-level assembly. We used the Chr19 sequence (NCBI AC No. WNKB01000034.1) from the previous version of human reference genome (CHM13v0.7) as input (Supplementary Fig. 4) and attempted to fill gaps with HiFi reads. Using DEGAP Ctglinker mode, the input sequences are automatically split into smaller contigs containing no gaps then get elongated by extending the edge sequences. As the result, two contiguous sequences were generated from nine fragmented contigs and the gap number was reduced from eight to one (Supplementary Fig. 4). The comparison result validates the high collinearity between DEGAP-resolved sequences and the gap-free CHM13v1.1 reference genome (Supplementary Fig. 4).

## Discussions

As more and more genomes are sequenced and assembled, gap-free genomes have gradually become the new standard for genome assembly. Assembling progresses from roughly getting longer N50 assembly operations to more detailed operations like resolve complex regions. DEGAP proves to be not only a valuable new tool assisting to generate a complete gap-free genome with PacBio HiFi reads, but also a powerful tool to aid in the resolution of complex regions. The biggest strength of DEGAP is that it dynamically adjusts the elongation strategy according to the complexity of the gap region and accurately elongates sequences even at low sequencing depth.

Accuracy and completeness are always the most important needs for a genome assembly, which may highly be affected by the quality and complexity of input raw data, especially in complex regions such as centromeres. DEGAP can distinguish HiFi reads from different sequences of high similarity, which allows it to resolve repetitive and complex regions in genomes, such as centromeric sequences. Understanding of the unique read depth value, reads accuracy, and the ratio of unique reads among all mapped reads may help us design DEGAP to dynamically recognize sequences in an elongation process.

For future genome assembly, we need to shift our focus from pursuing more complete genomes to complete and also precise genome. DEGAP can be used as a tool to verify the quality of genome assembly. In each round of extension steps, since all reads aligned to edge sequence are rigorously tested and evaluated, the extension results of DEGAP can also be used as a reference to determine the correctness of local assembly by finding the difficult-to-be-assembled genome regions then comparing the differences between DEGAP extension result and other assembly results. DEGAP, as a new ‘post’ genome assembly software tool, will play a powerful role that help to build better quality genome sequences of longer contiguity.

## Methods

### Dynamic HiFi reads finding

DEGAP uses minimap2^(8)^ to generate the sequence alignment file in SAM/BAM format, sets the distance from the boundary threshold from 10 bp to 500 bp (the default is 500 bp and can be customized with the parameter “--edge”), MAPQ from 20 to 0, alignment length from 3,000 bp to 500 bp and extension length from 1,000 bp to 10 bp. After selecting all aligned HiFi reads and getting the extension HiFi reads in a loose threshold, DEGAP adaptively adjusts the filtering thresholds from a stringent one for the most suitable HiFi reads (Supplement Fig. 1). To reduce the screening time, DEGAP selects the entire HiFi reads with parameters as NM (Edit distance to the reference, including ambiguous bases but excluding clipping)/Alignment length <0.1, MAPQ >=0, alignment length >=500, extension length >=10 and distance from boundary <=500. If no extension reads found, DEGAP stops the entire elongation process and exports the final result. In this step, the alignment file was processed by SAMtools^(13)^ and pysam python package.

### Dynamic extension sequence finding

DEGAP assembles the extension reads by hifiasm^(6)^ and aligns the assembled sequences to edge sequences by MUMmer^(14)^. The minimum alignment length may be set from 1000 bp to 500 bp, minimum alignment identity from 99.0 to 95.0, distance from boundary from 10 bp to maximum extension length. If there are no sequence assembled by hifiasm pass the selection process, DEGAP uses the common sequences between any two HiFi reads (DEGAP uses the entire sequence if only one extension read found). With the common sequences, DEGAP runs the dynamic selection process again to find the best extension sequence. The goal of this process is to find the best extension sequence by selecting the alignment blocks from hifiasm assembly or common sequences, which may contain multiple sequences (Supplement Fig. 3a).

### Fuzzy Alignment generation

DEGAP uses MUMmer to align the assembly or common sequences with the edge sequence to find the best alignment result which can elongate the edge sequence (Supplement Fig. 3a, b). DEGAP uses the length of the alignment area, alignment identity and distance from boundary to select the alignment result, which is similar to the extension reads selection. And if the alignment is reversed, DEGAP takes the reverse complement of the extension sequence (Supplement Fig. 3b). If there are multiple alignment results, DEGAP chooses the most solid alignment block (Supplement Fig. 3c). Moreover, DEGAP also considers some special circumstance that hifiasm’s assembly cannot elongate the edge sequence.

In case of the two sequences are greatly different or contain tandem repeats, the result does not always show a single alignment block at the edge (Supplementary Fig. 3d, e). When getting more than one alignment block but none can pass the dynamic selection, DEGAP generates fuzzy alignment result to test if the extension sequence can be utilized (Supplement Fig. 3d, e).

## Data availability

All described datasets are publicly available. MH63RS3 raw sequencing data used for this project are archived at NCBI under accession SRX6957825, SRX6908794. The genome assemblies are available at NCBI under CP054676–CP054688 or at the National Genomics Data Center under BioProject no. PRJCA005549. Human reference genome GRCh38.p13 sequence data and genome browser are available from https://github.com/marbl/CHM13.

## Code availability

DEGAP was developed as an open-source tool on a Linux platform with the Python programming language and could be downloaded at https://github.com/Jianwei-Zhang/DEGAP.

