## Supplementary Information for "DEGAP: Dynamic Elongation of a Genome Assembly Path"

### Supplementary Figures

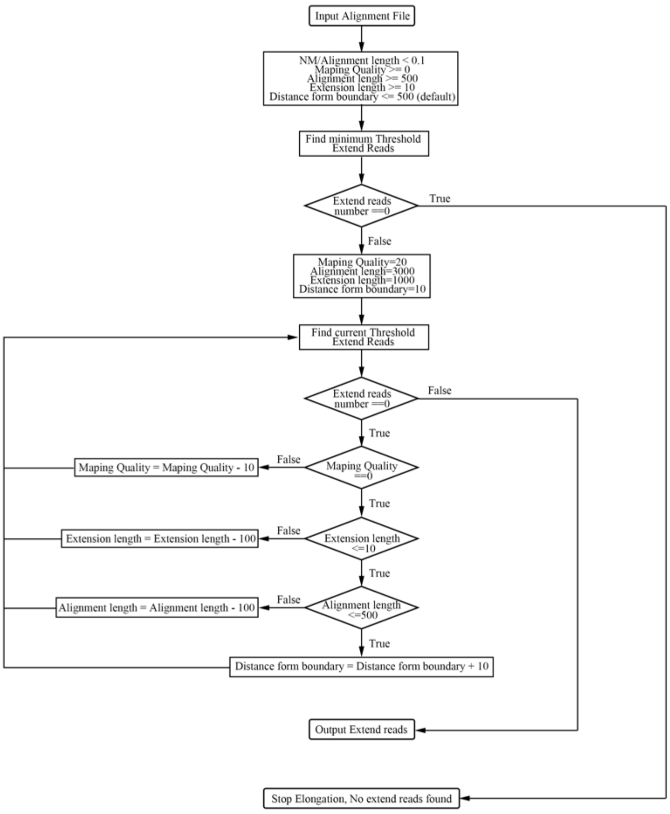


**Supplementary Figure 1:** Pipeline of dynamic HiFi reads finding.

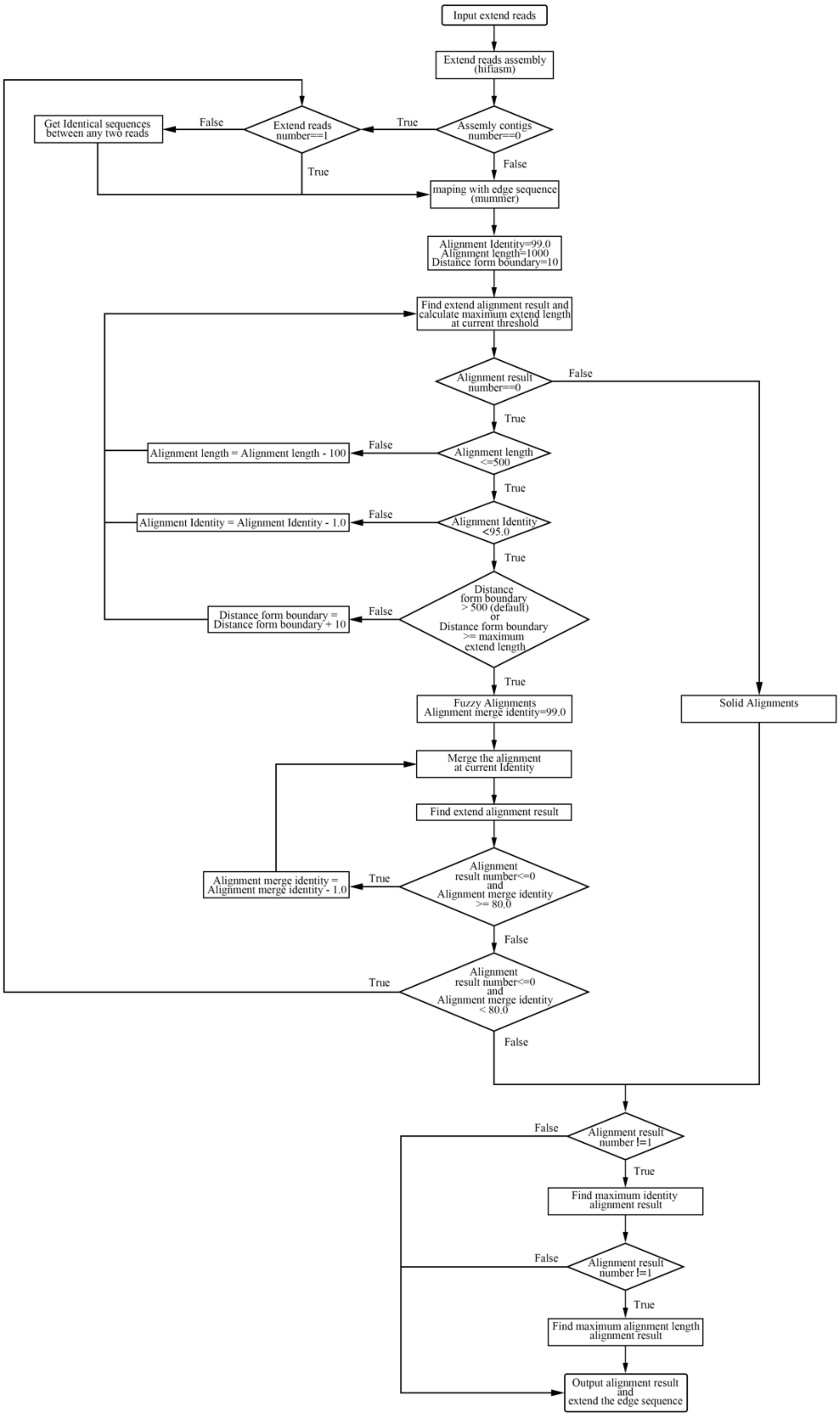


**Supplementary Figure 2:** Pipeline of dynamic extension sequence finding.

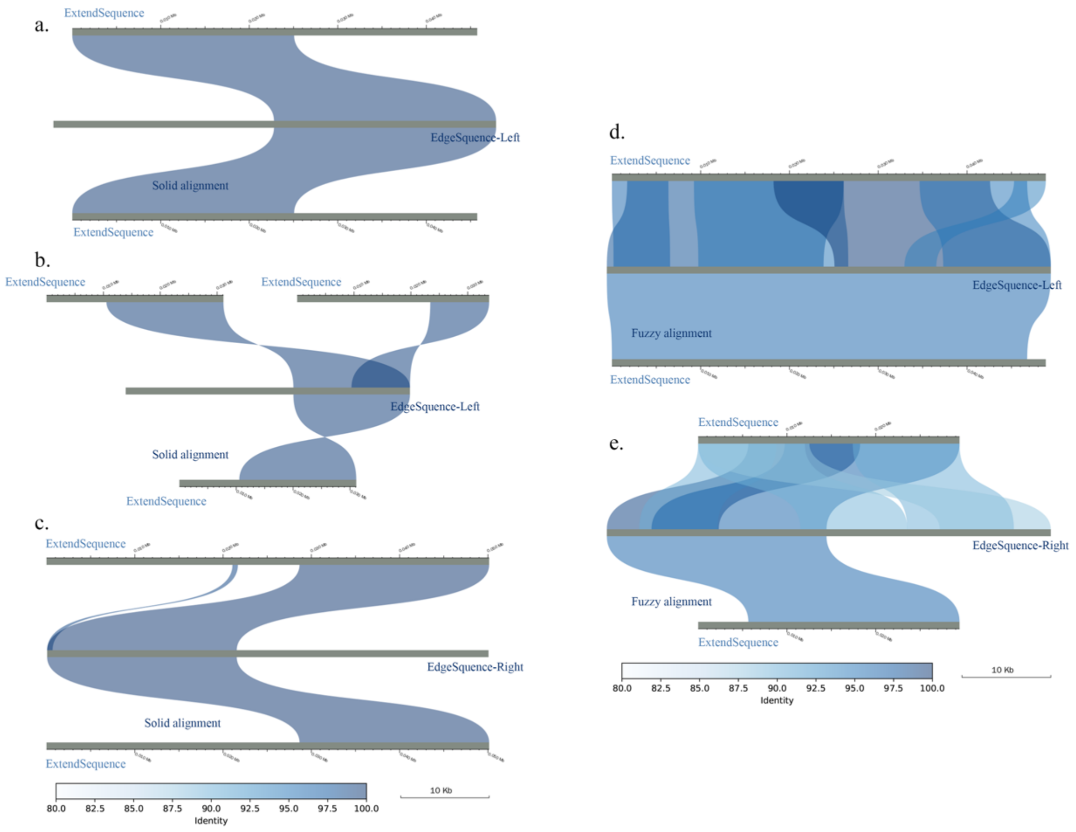


**Supplementary Figure 3: Solid alignment selection and fuzzy alignment generation. a**, Solid alignment selection. From top to bottom are the input sequence alignment, edge sequence and alignment via dynamic selection. **b**, Dynamic extension sequence selection. DEGAP selects the best extend sequence from multiple potential extend sequence. **c**, DEGAP remove alignments that are not useful for elongating sequences. **d**, Fuzzy alignment generation. From top to bottom are raw alignment result and the fuzzy alignment result. **e**, Removal of useless alignments and generation of fuzzy alignment.

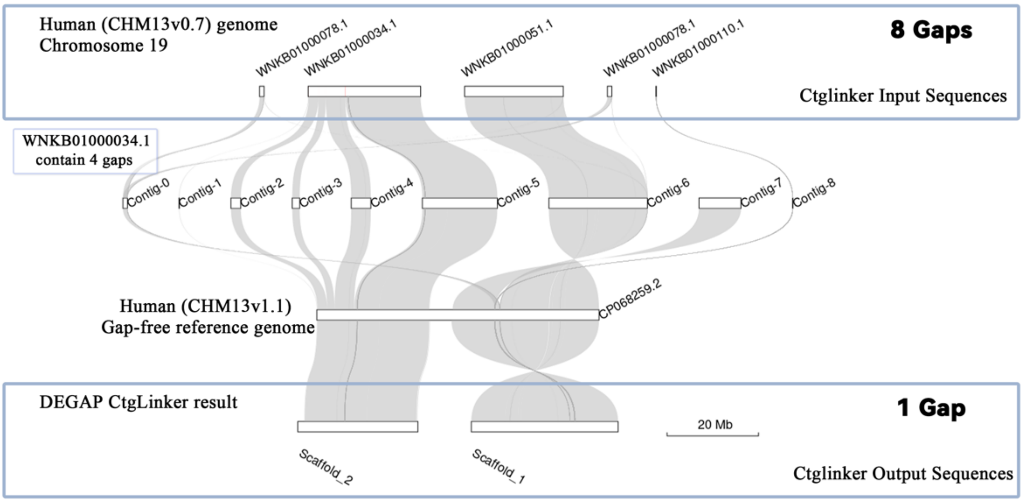


**Supplementary Figure 4: DEGAP for Human (CHM13v0.7) Chromosome in CtgLinker mode.** DEGAP ordered and linked the contigs properly and reduce gap number from eight to one. From top to bottom are the human (CHM13v0.7) Chr19 sequence (DAGAP input sequences), Contigs separated from the human (CHM13v0.7) Chr19 sequence by gaps, the human (CHM13v1.1) Chr19 gap-free reference sequence, and the DEGAP output sequences.
